# Waning and boosting of functional humoral immunity to SARS-CoV-2

**DOI:** 10.1101/2022.07.22.501163

**Authors:** X. Tong, R.P. McNamara, M.J. Avendaño, E.F. Serrano, T. García-Salum, C. Pardo-Roa, J. Levican, E. Poblete, E. Salina, A. Muñoz, A. Riquelme, G. Alter, R.A. Medina

## Abstract

Since the emergence of the SARS-CoV-2 virus, we have witnessed a revolution in vaccine development with the rapid emergence and deployment of both traditional and novel vaccine platforms. The inactivated CoronaVac vaccine and the mRNA-based Pfizer/BNT162b2 vaccine are among the most widely distributed vaccines, both demonstrating high, albeit variable, vaccine effectiveness against severe COVID-19 over time. Beyond the ability of the vaccines to generate neutralizing antibodies, antibodies can attenuate disease via their ability to recruit the cytotoxic and opsinophagocytic functions of the immune response. However, whether Fc-effector functions are induced differentially, wane with different kinetics, and are boostable, remains unknown. Here, using systems serology, we profiled the Fc-effector profiles induced by the CoronaVac and BNT162b2 vaccines, over time. Despite the significantly higher antibody functional responses induced by the BNT162b2 vaccine, CoronaVac responses waned more slowly, albeit still found at levels below those present in the systemic circulation of BNT162b2 immunized individuals. However, mRNA boosting of the CoronaVac vaccine responses resulted in the induction of significantly higher peak antibody functional responses with increased humoral breadth, including to Omicron. Collectively, the data presented here point to striking differences in vaccine platform-induced functional humoral immune responses, that wane with different kinetics, and can be functionally rescued and expanded with boosting.

## Introduction

Severe acute respiratory syndrome coronavirus-2 (SARS-CoV-2) is the causative agent of coronavirus disease-2019 (COVID-19). Since it was first identified in late 2019 (1–3), more than half a dozen vaccines, using novel and diverse platforms, have been deployed globally to provide protection against this highly transmissible pathogen (4). However, despite the remarkable success of the vaccines, the virus has undergone adaptations that have facilitated transmission among humans, and with mutations selectively accumulating in the receptor-binding domain (RBD), permitting the virus to also escape vaccine-induced neutralizing antibody responses (5)(6–8). In fact, several variants of concern (VOC) have arisen throughout the world since the onset of the pandemic, causing selective waves of transmission (9–14), most strikingly observed with the emergence of the Omicron VOC that has led to remarkable global spread (15). Despite the fact that these emerging VOCs can subvert neutralization and spread with extreme ease, severe disease and death have not increased proportionally to the spread of the disease. Instead, in unvaccinated populations, all VOCs, including Omicron, have led to severe disease and death (16–18), arguing that the vaccine-induced non-neutralizing immune responses are key to attenuating disease.

Humoral immunity, including both binding and neutralizing antibody titers, has been tightly linked to protection against COVID-19 in phase 3 vaccine trials (19–22). However, beyond the ability of antibodies to bind and block infection, binding antibodies also can leverage the innate immune system to capture, kill, and clear viruses or infected cells via their ability to interact with Fc-receptors present on all immune cells (23–25). These non-neutralizing antibody functions selectively evolve in individuals that survive severe disease (26), are associated with the therapeutic activity of convalescent plasma therapy (27), are associated with vaccine-mediated protection in the non-human primate model (28), and contribute to the therapeutic activity of the monoclonal antibodies (29). Moreover, recent data suggest that even vaccines using the same technology (i.e., mRNA) can elicit significantly different functional humoral immune responses (24). Nonetheless, while antibody titers and neutralization wane significantly across vaccine platforms, it is unclear whether functional immunity wanes concomitantly to titers and/or whether the waned immunity can be boosted efficiently.

Among the vaccines that have been deployed globally, the inactivated CoronaVac (Sinovac) vaccine and the mRNA Pfizer/BioNTech BNT162b2 vaccine have been two of the most broadly deployed vaccines globally, having been administered to billions of individuals. In phase 3 trials, the CoronaVac vaccine exhibited 84% protection against severe COVID-19 disease, whereas the BNT162b2 vaccine exhibited 95% protection (30–32). Differences in antibody and neutralizing titers across the vaccine platforms have been proposed as critical determinants of different efficacy (33). However, whether these platforms raise distinct overall functional humoral immune responses if these wane at similar rates, and whether CoronaVac immunity can be augmented, potentially in the setting of an mRNA-vaccine boost, remains unclear.

Here we deeply profiled the functional humoral immune response induced by CoronaVac and Pfizer/BNT162b2 vaccines, particularly how the vaccine-induced functional responses waned with time and analyzed how boosting with the BNT162b2 vaccine restored the waned CoronaVac immunity. We identified that Fc-receptor binding antibodies waned much faster than binding IgG1. Moreover, while antibody functions were induced to Omicron following BNT162b2 vaccination, these responses waned rapidly over time and were not observed in the setting of CoronaVac vaccination. Importantly, mRNA boosting of the CoronaVac response yielded a striking enhancement of functional humoral immunity across VOCs, including Omicron. These data collectively point to striking vaccine platform-based differences in peak functional humoral protection, as well as their distinct waning profiles. This phenotype can be reversed and, in several cases, functionally expanded by heterologous boosting.

## Results

### Inactivated and mRNA COVID-19 vaccines induce spike- and RBD-specific antibodies that wane with time

Beyond their ability to neutralize infection, antibodies can attenuate disease via their ability to bind to virus or virally infected cells and then recruit the innate immune system at the site of infection, via Fc-receptors (24, 25, 27, 28). As a result, the persistence and breadth of binding of Spike-specific antibodies is a key initial determinant of potential non-neutralizing vaccine effectiveness. Thus, we employed a systems serology approach to deeply profile the peak immunogenicity and decay profiles of SARS-CoV-2- and VOC-specific antibodies following CoronaVac and Pfizer/BNT162b2 vaccination (**Figure 1A**). Two doses of both vaccines elicited detectable IgG1 against WT SARS-CoV-2 Spike and Alpha and Beta VOC (**Figure 1B,** left) that waned over time resulting in more similar titers across the vaccine platforms at 4-5 months following immunization compared to peak immunogenicity. This dose-dependent increase and subsequent waning of receptor-binding domain (RBD) antibodies followed a similar trajectory (**Figure 1B**, right). Other VOCs such as Gamma and Delta displayed similar patterns for IgG1 recognition against full-length Spike or RBD after vaccination with BNT162b2 or CoronaVac (**Figure 1C**). This phenotype was specific to SARS-CoV-2 VOC (Supplementary Figure 1). As expected, CoronaVac or BNT162b2 vaccination drove specific IgG1 responses against most VOCs, which also waned with time. Notably, Omicron Spike and RBD IgG1 responses were lower than responses observed for other VOCs in BNT162b2 and nearly undetectable in most CoronaVac recipients (**Figure 1C,** bottom row).

**Figure 1.**
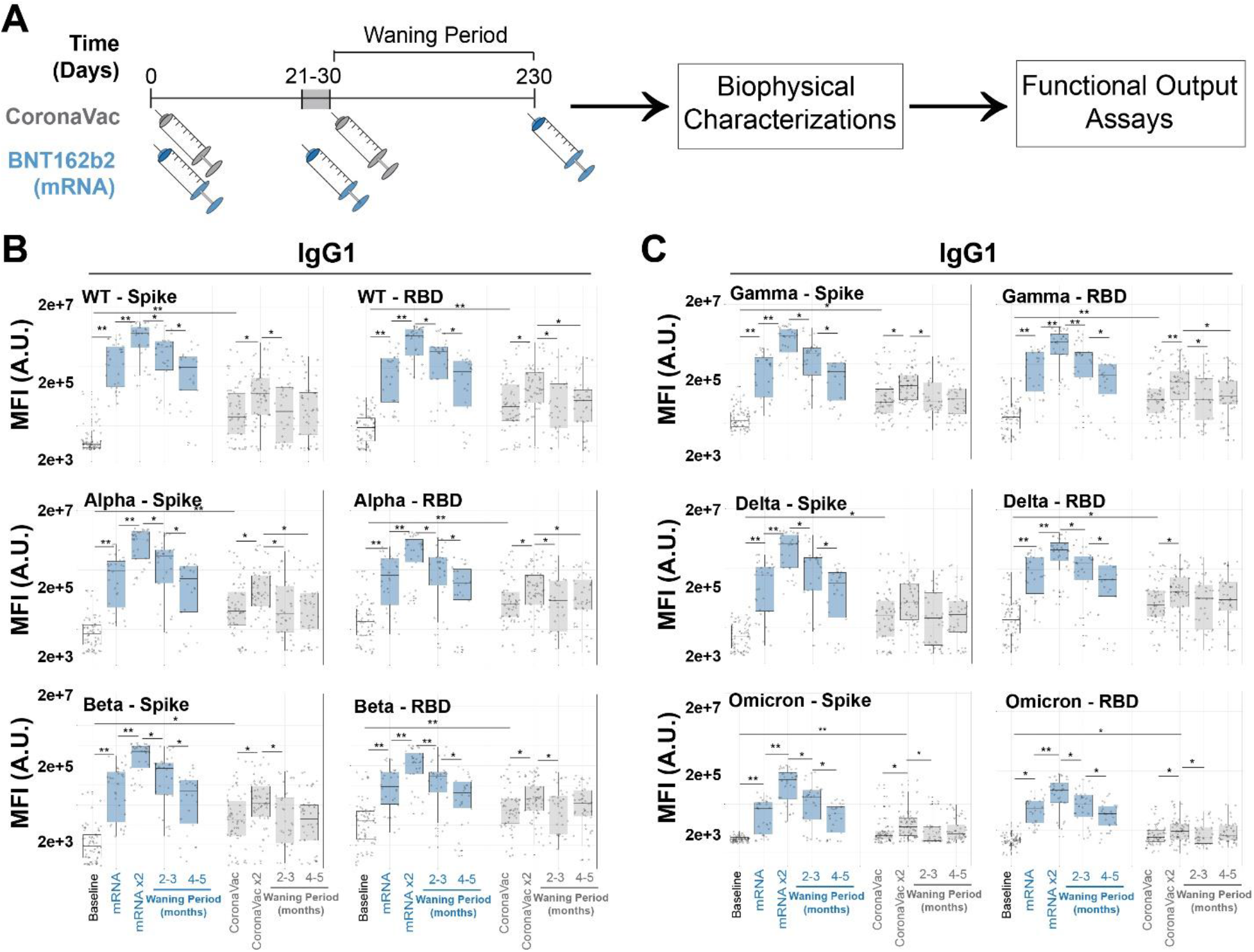
COVID-19 vaccines exhibit a pan-variant waning of Spike-binding antibodies. (A) Scheme for vaccination series and subsequent systems serology analysis. Subjects who received two doses of the inactivated vaccine CoronaVac were boosted with the mRNA vaccine BNT162b2. As controls, the two-dose mRNA vaccine recipients were also analyzed. Sera were collected at various time points and systems serology assays were conducted. (B) Spike- (left) and receptor binding domain (RBD) (right) specific IgG1 levels for SARS-CoV-2 WT and early VOCs (Alpha and Beta) were quantified in the baseline (prior to immunization, white, column 1), 1- and 2-dose BNT162b2 mRNA (blue, columns 2 and 3, waning periods in lanes 4 and 5), 1- and 2-dose CoronaVac (gray, columns 6 and 7, waning periods in lanes 8 and 9) via Luminex systems serology. Y-axis represents the MFI of binding in arbitrary units (A.U.) of a specific antigen. Shown are box and whiskers, along with individual data points, which represent the mean of technical replicates on individual patients. (C) Same as B, but for latter VOC (Gamma, Delta, and Omicron) of SARS-CoV-2 VOC. * = p < 0.05 and ** = p < 0.01, Kruskal-Wallis test, for all panels.

### Inactivated and mRNA COVID-19 vaccines exhibit distinct Fc-receptor waning

To drive antibody-effector functions, antibodies must interact with Fc-receptors found on all innate immune cells (25, 34, 35). Thus we next assessed whether vaccine-induced FcR binding profiles, specifically the binding to the 4 human low-affinity Fcγ-receptors (FcγR2A, FcγR2B, FcγR3A, FcγR3B), waned with similar or disparate kinetics to antibody titers across the vaccine platforms. We observed distinct mRNA vaccine induced FcR binding maturation and decay across the FcRs. Specifically, opsinophagocytic activating FcγR2A-binding Spike specific antibodies emerged rapidly after the first dose, peaking after the second dose, and remained detectable 4-5 months after vaccination, despite decaying to levels lower than observed after the first mRNA vaccine dose (**Figure 2A**). Conversely, FcγR2B, FcγR3A, and FcγR3B-binding Spike-specific levels were induced more slowly after the first mRNA-vaccine dose, requiring the second dose to mature fully (**Figure 2B-D**). These responses then declined rapidly over time, with nearly undetectable levels in most vaccinees 4-5 months following the completion of the primary mRNA vaccine series of immunization. These data point, for the first time, differentiated waning across FcR-binding antibodies following mRNA vaccination, with the more robust persistence of FcγR2A-binding Spike-specific antibodies over time.

**Figure 2.**
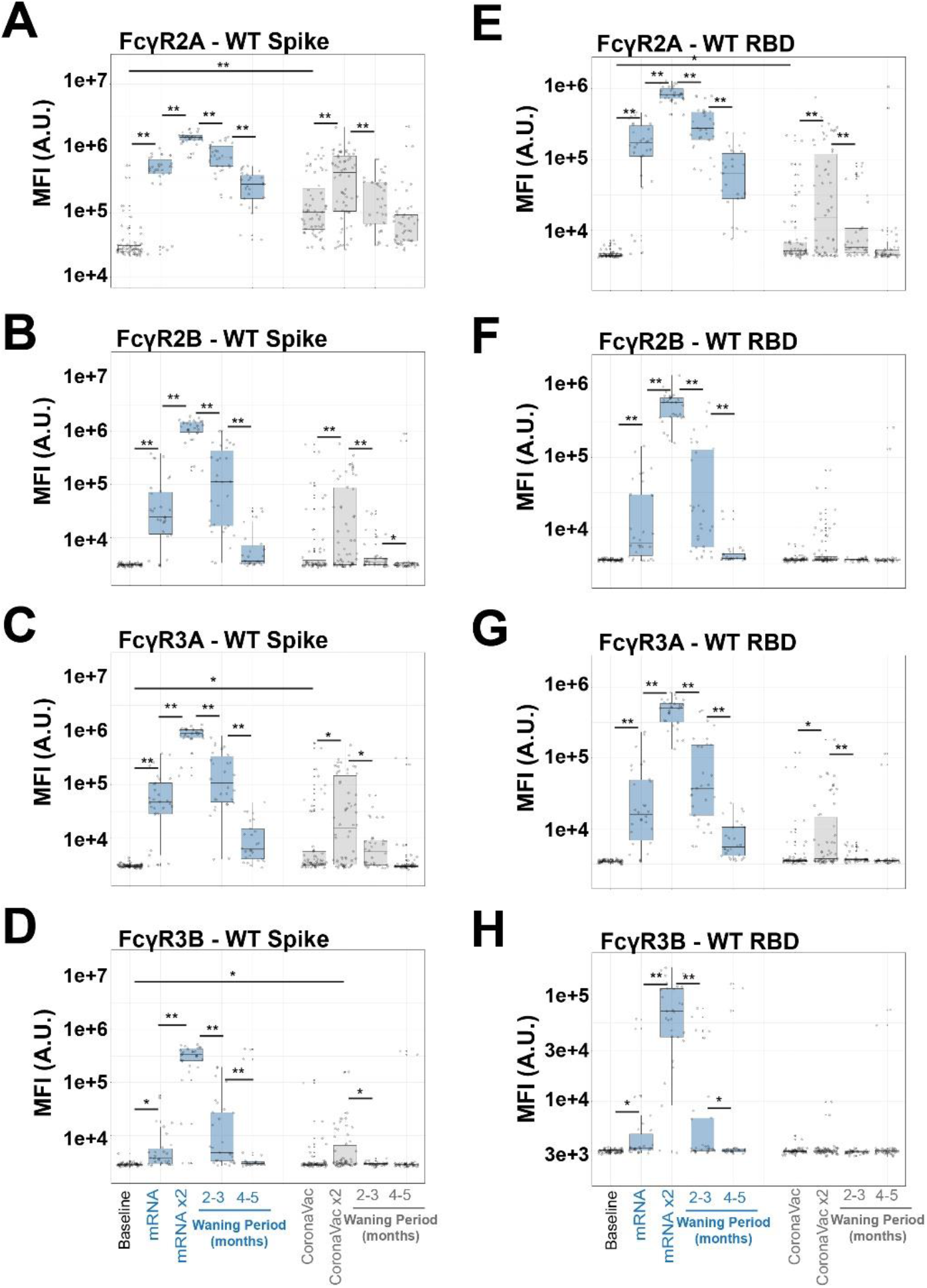
FcR binding antibodies towards full-length Spike and RBD have rapid wane kinetics. (A-D) FcR antibody binding towards WT SARS-CoV-2 Spike was quantified in the baseline (prior to immunization, white, column 1), 1- and 2-dose BNT162b2 mRNA (blue, columns 2 and 3, waning periods in lanes 4 and 5), 1- and 2-dose CoronaVac (gray, columns 6 and 7, waning periods in lanes 8 and 9) via Luminex systems serology. Y-axis represents the MFI of binding full-length WT SARS-CoV-2 Spike. Shown are box and whiskers, along with individual data points which represent the mean of technical replicates on individual patients on (A) FcγR2A, (B) FcγR2B, (C) FcγR3A, and (D) FcγR3B. (E-H) Same as A-D, but for SARS-CoV-2 RBD. * = p < 0.05 and ** = p < 0.01, Kruskal-Wallis test for all panels. Note that Y-axis scales are not the same. See also Supplementary Figure 1.

Despite the clear induction of binding IgG antibodies by the CoronaVac vaccine, FcR binding profiles exhibited significant differences, both in peak titer and in waning kinetics. Opsinophagocytic FcγR2A and cytotoxic FcγR3A binding antibodies were induced most strongly by this vaccine in a dose-dependent manner. Strikingly, a single dose of the vaccine-induced low levels of these FcR binding antibodies. FcγR2A-binding Spike-specific antibodies accumulated after the second dose of the vaccine but declined to near background levels 4-5 months following completion of the primary 2 dose series of immunizations. A subset of vaccinees further induced activating cytotoxic FcγR3A binding antibodies after the second dose of the vaccine, but all waned to undetectable levels. No inhibitory FcγR2B or neutrophil-specific FcγR3B-binding antibodies were induced by the first dose of the CoronaVac vaccine. Second-dose responses were present, but these quickly waned to background levels across all vaccinees (**Figure 2B and D**) despite the clear induction of binding IgG antibodies (**Figure 1**).

The same profiling was performed for RBD-specific antibodies across the 2 vaccine platforms. Again, BNT162b2 mRNA vaccination induced FcγR2A-binding RBD-specific antibodies after a single dose, which matured exponentially after a second dose (**Figure 2E**). However, these RBD-specific FcγR2A-binding antibodies decayed rapidly over 4-5 months. Conversely, very low, although detectable, levels of RBD-specific FcγR2A-binding antibodies were induced by the CoronaVac vaccine that fully decayed over 4-5 months (**Figure 2E**). Interestingly, 2 doses of BNT162b2 mRNA vaccination were able to drive RBD-specific antibodies capable to interact with FcγR2B and FcγR3B binding antibodies, whereas a single dose produced low and heterogenous responses (**Figure 2F-H**). Unlike FcγR2A-binding antibodies, these waned to low-to-undetectable levels over 4-5 months. In contrast, the CoronaVac vaccination did not induce any RBD-binding antibodies able to interact with FcγR2B or FcγR3B and only minimally produced FcγR3A binding antibodies, and these waned after 3 months. These data argue that both vaccines induced Spike and RBD-specific IgG1, despite showing distinct functional FcR binding properties. These functional properties waned more rapidly compared to the binding antibodies.

### mRNA vaccine boosts antibody and Fc effector functions to recognize SARS-CoV-2 VOC

Given the comparatively low antibody levels induced by the CoronaVac vaccine against all VOCs, particularly Omicron, we next investigated whether mRNA boosting of previous CoronaVac vaccinees could augment antibody breadth and Fc-effector function. Upon mRNA boosting, a sharp and pan-VOC increase in IgG1 levels was observed for full-length Spike (**Figure 3A**) and for the RBD of all VOCs (**Figure 3B**). This was particularly striking for Omicron-specific IgG responses, which exhibited the lowest cross-reactive IgG1 levels prior to boosting, yet Omicron-specific immunity was boosted to similar levels to other VOC Spike-specific responses (**Figure 3A**), albeit Omicron RBD recognition was boosted but did not reach the levels of other RBD VOC responses.

**Figure 3.**
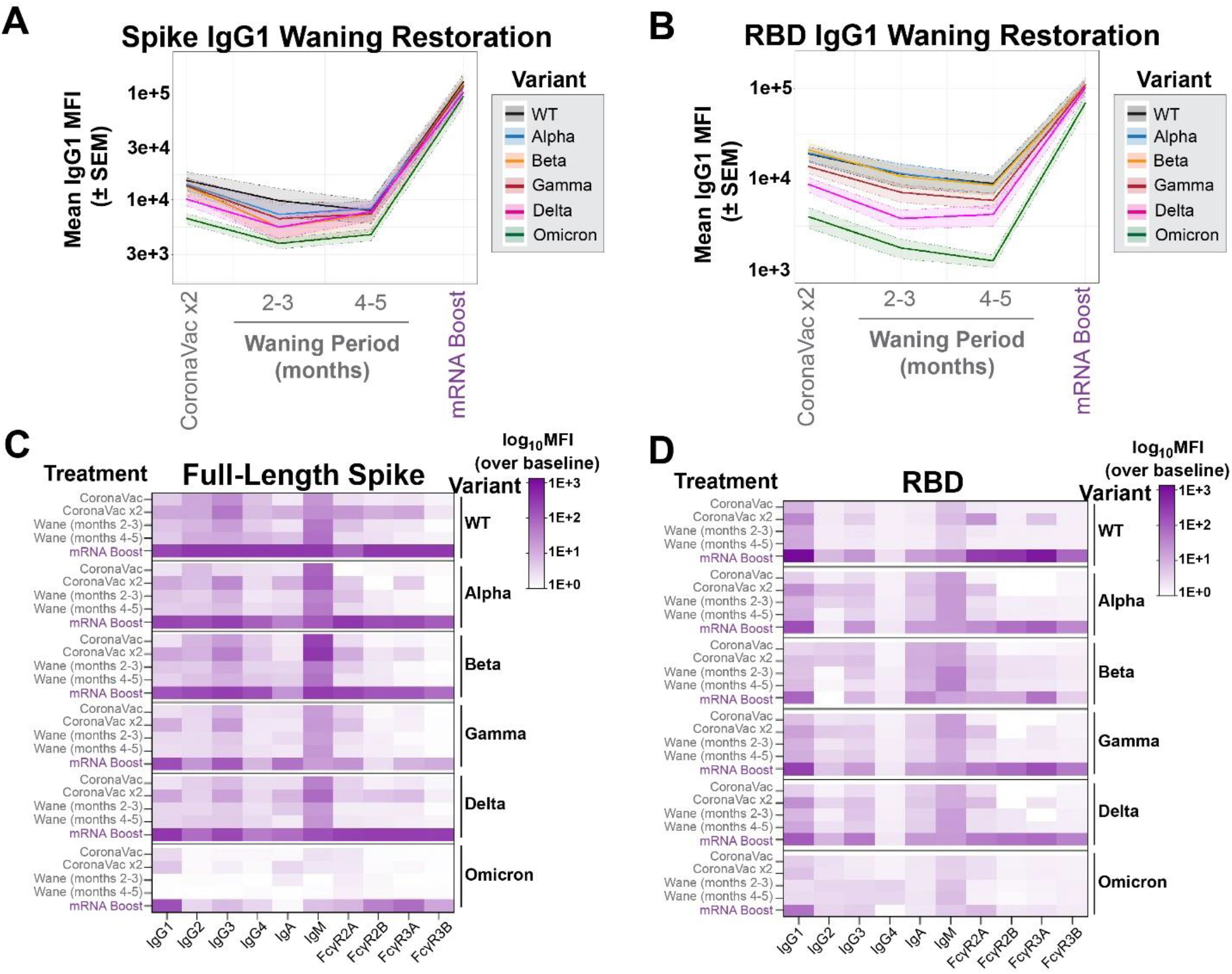
COVID-19 mRNA boosters can enhance responses of multiple Spike-specific antibody-subclasses and -isotypes, and functional Fcγ-receptor complexes. (A) Spike-specific IgG1 levels were measured at peak immunogenicity following the two-dose CoronaVac vaccination series (lane 1), during waning periods (columns 2 and 3), and after mRNA boost (column 4) for SARS-CoV-2 VOC (color legend shown on right). The Y-axis represents the MFI of the Spike from the VOC. Shown are means (solid line) with SEM for each VOC in the corresponding color in the shaded region. (B) Same as A, but for the receptor-binding domains (RBD) of SARS-CoV-2 VOC. (C) Heatmap representation of binding for full-length Spike by antibody-subclasses and -isotypes and Fcγ-receptor complexes. Shown on the left are the treatments of CoronaVac doses, the subsequent waning period, and mRNA-vaccine booster, and on the right are the SARS-CoV-2 VOC or WT. The scale bar is shown to the right of the heatmap and represents MFI over baseline values. The values in each region represent the mean. (D) Same as C, but for the RBD of WT SARS-CoV-2 and VOC. See also Supplementary Figure 2.

To further examine the overall antibody isotype, subclass, and FcR binding breadth, we next examined the dynamics of the overall humoral profile of CoronaVac-induced SARS-CoV-2 specific antibodies over time and after boosting (**Figure 3C**). As expected, WT Spike-specific response had the highest and most consistent initial response but waned over the course of 5 months. This waning effect was not only rescued but significantly boosted over peak titers following the Pfizer/BNT162b2 dose. This boost also raised highly functional IgG3- and IgM-responses to Spike, pointing to the induction of new responses. IgA responses evolved over time following CoronaVac immunization and were robustly boosted by the BNT162b2 response. Conversely, FcR-binding antibodies, key for eliciting antibody effector functions, were observed after the second CoronaVac dose and then quickly returned to near undetectable levels. However, following the BNT162b2 booster, all Spike-specific FcR binding responses were robustly enhanced against the WT Spike antigen.

Similar trends in subclass and isotype profiles were observed across VOCs. However, notably, Spike-specific IgA responses were less significantly induced to the Beta and Gamma-variants, potentially accounting for some potential mucosal transmission liabilities for this VOC. Additionally, IgG2 and IgG4 responses were induced weakly to the Gamma Spike. Omicron Spike-specific binding antibody FcR profiles differed most across the VOCs, with limited subclass and isotype responses to the Omicron Spike, despite robust IgG1 responses (**Figure 3C**). Moreover, weaker opsinophagocytic FcγR2A-binding and FcγR3B-binding Spike-specific binding antibodies were noted following the BNT162b2 boosting. In contrast, robust FcγR2B and FcγR3A binding antibodies were induced, pointing to the selective induction of individual FcR binding responses to Omicron following boosting.

The cross-VOC RBD-specific response was more variable (**Figure 3D**). While RBD-specific IgG1 responses were detected at nearly all time points following CoronaVac vaccination, BNT162b2 boosting resulted in a robust augmentation of RBD-specific IgG1 responses, superior to those observed at peak immunogenicity. Limited subclass and isotype responses were observed to the wildtype RBD with CoronaVac immunization alone. However, the addition of the mRNA booster significantly raised RBD-specific IgG3, IgA, IgM, and all FcR binding responses. A similar profile was observed to the Alpha, Gamma, and Delta VOC RBDs. However, the mRNA vaccine boosts only induced cross-reactive Beta RBD-specific antibodies with a more limited FcR binding profile, and a preferential significantly higher FcγR3A binding profile. Interestingly, while cross-reactive Omicron RBD-specific IgG1 were induced with the BTN162b2 boost, the boost failed to induce Omicron RBD-specific antibodies of distinct subclasses, isotypes or with broad FcR binding profiles. These data suggest that mRNA boosting can broaden the subclass, isotype, and FcR binding profile across VOCs (Supplementary Figure 2), but may only partially rescue FcR binding specifically to more distant VOCs, particularly to those that have spread more efficiently and to which CoronaVac was shown to be slightly less efficient (36).

### mRNA-Vaccine boosting of CoronaVac recipients restores and broadens functional humoral defenses

While FcR binding is required to initiate Fc-effector function, we next sought to profile the longitudinal functional vaccine-induced humoral immune responses across the vaccine platforms, focused on the WT and Omicron SARS-CoV-2 Spike antigens. Although the CoronaVac vaccine elicited a moderate level of WT Spike-specific complement deposition (ADCD) (grey line) after the primary vaccine series, it quickly waned (64% and 72% reduction from peak activity at 2-3 months and 4-5 months from peak activity, respectively). The BNT162b2 vaccination elicited highly robust ADCD responses, that persisted for the first 2-3 months following the primary immunization series, but then the response waned at 4-5 months to a 58% overall reduction from peak activity (**Figure 4A**). However, BNT162b2 boosting of CoronaVac vaccinees resulted in a highly significant expansion of ADCD responses, to levels higher than peak BNT162b2 mRNA-induced responses alone. Conversely, neither vaccine series elicited comparable immunity to the Omicron Spike (purple and blue lines on the left and right panels, respectively), although mRNA-vaccine boosting was noted.

**Figure 4.**
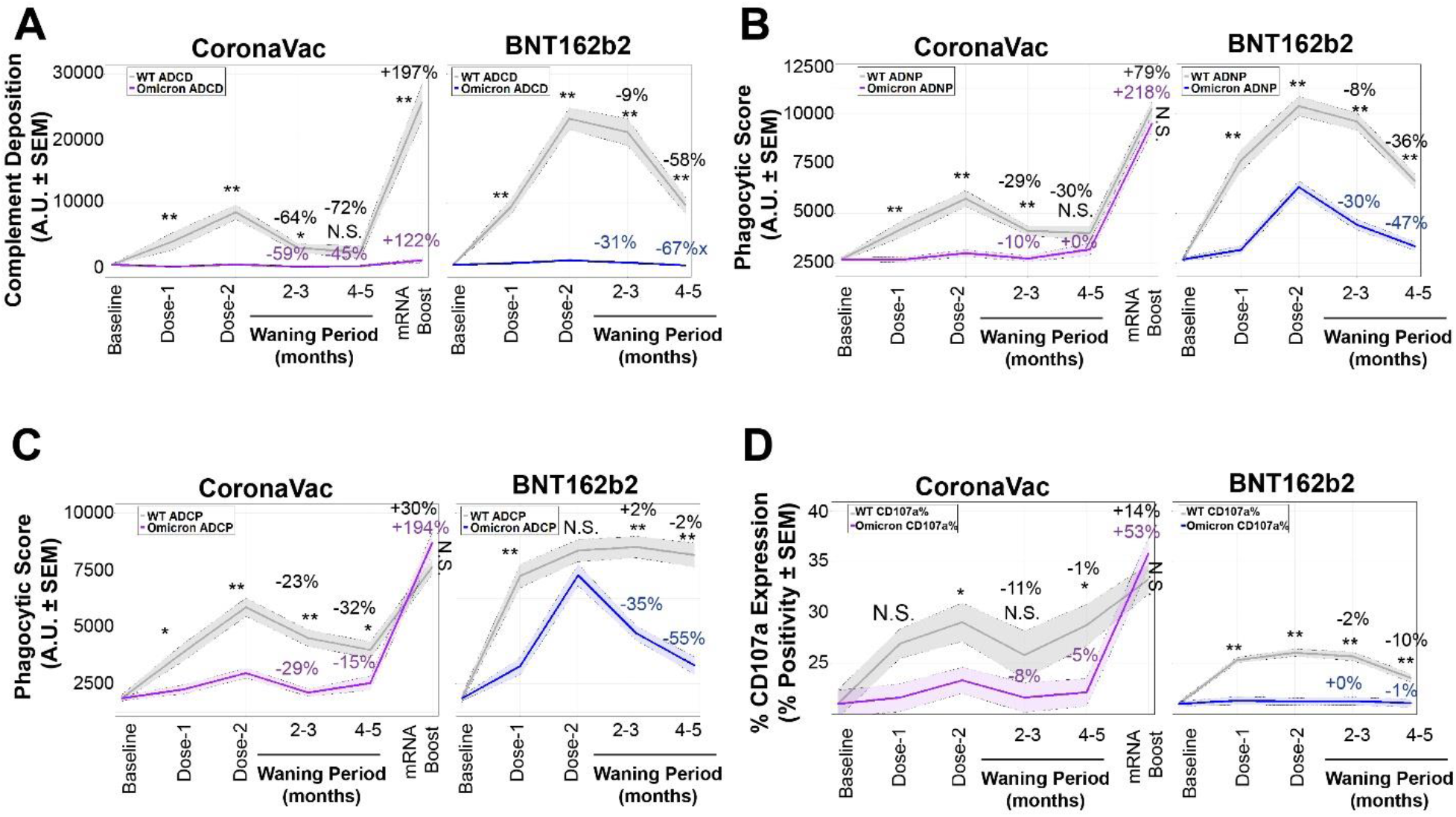
mRNA-vaccine boosting of CoronaVac recipients broadens humoral defenses, including towards Omicron. (A) (Left) Antibody-dependent complement deposition (ADCD) measured in fluorescent arbitrary units (A.U.) in individuals at baseline (column 1) who received two doses of CoronaVac (columns 2 and 3), during the waning period (columns 4 and 5), and after mRNA-vaccine booster (column 6); WT Spike is shown in gray and Omicron is shown in purple. Peak functionality was set to responses after dose 2, and percent changes during the waning and boosting are shown above the timepoints. (Right) ADCD against WT Spike (gray) and Omicron Spike (blue) in individuals at baseline (column 1), after two doses of an mRNA vaccine (columns 2 and 3), and during the waning period (columns 4 and 5). Peak functionality was set to responses after dose 2, and percent changes during the waning and boosting are shown above the timepoints. Shown are the mean (solid line) and standard errors of the mean (SEM, shaded areas). (B) Same as A, but for antibody-dependent neutrophil phagocytosis (ADNP) measured in phagocytic A.U. (C) Same as A but for antibody-dependent monocytic cellular phagocytosis (ADCP) measured in phagocytic A.U. (D) Primary natural killer (NK) cells were measured for percent surface expression of CD107a after incubation with sera from (left) CoronaVac + mRNA-boosted individuals or (right) two-dose mRNA vaccine recipients for WT and Omicron Spike. * = p < 0.05, ** = p < 0.01, and N.S. = not significant for all WT and Omicron comparisons (paired T-tests). See also Supplementary Figure 3.

Similarly, WT Spike-specific antibody-dependent neutrophil phagocytosis (ADNP) was induced by both vaccines, albeit to lower levels with CoronaVac, although ADNP activities waned 30% and 36% after 4-5 months in CoronaVac and BNT162b2 recipients, respectively (**Figure 4B**). However, after BNT162b2 boosting, ADNP levels in CoronaVac vaccinees rose to comparable to those observed after the primary series with the BNT162b2 vaccine. Importantly, the low-level Omicron-specific ADNP activity was noted after the second dose of the CoronaVac primary series, which declined rapidly. BNT162b2 boosting led to robust induction of Omicron-specific ADNP activity that was similar to WT Spike.

A nearly identical pattern was observed with antibody-dependent monocyte cellular phagocytosis (ADCP), albeit ADCP did not decline after the primary series of BNT162b2 vaccination for WT Spike (**Figure 4C**). Again, while low Omicron ADCP was induced by the CoronaVac vaccine that waned, a cross-reactive response was restored with the BNT162b2 immunization, which is comparable to the WT-specific primary ADCP levels induced by the boost. Finally, WT-specific NK cell activation measured by NK cell degranulation through percent CD107a positivity (**Figure 4D**) or macrophage-inflammatory protein 1b (MIP-1b) expression (Supplementary Figure 3), was surprisingly elicited to comparable levels across CoronaVac and BNT162b2 vaccination after the primary series to the WT Spike antigens. Omicron-specific NK cell activating antibodies were only minimally observed among CoronaVac vaccinees after the primary series. However, after the BNT162b2 boost, both WT and Omicron-specific NK cell activating antibodies were significantly upregulated despite no activity being detected in the BNT162b2 recipients (**Figure 4D**, right), pointing to a distinct functional maturation in the pre-existing setting of the CoronaVac recipients.

## Discussion

Emerging data suggest that distinct COVID-19 vaccination platforms can establish humoral responses with varying breath and magnitude towards Spike VOCs (4, 24, 36). However, these responses wane rapidly across vaccine platforms (20, 37). Yet in addition to binding and neutralization, vaccine-induced antibodies are also capable of mediating an array of antibody effector functions (23, 28), that have been linked to protection against severe disease and death (26) and monoclonal therapeutic activity (25, 38). However, whether these responses decay at the same rate as neutralization remains incompletely understood. A comparison of the 2 most widely distributed global COVID-19 vaccines, the CoronaVac and Pfizer/BioNTech mRNA vaccines revealed striking differences in peak antibody binding titers and Fc-effector functions across the 2 vaccine platforms, that both waned with time. Moreover, CoronaVac-induced Fc-effector functions demonstrated a steep and rapid decline to undetectable levels while binding to IgG was still present. These data suggest that the breadth of antibody binding does not directly translate to the level of circulating antibody effector function. Yet critically, a Pfizer/BioNTech boost of CoronaVac immunity restored antibody effector function, in some cases above those observed with mRNA immunization alone. While the longitudinal duration of these boosted functional responses remains unclear, our results clearly show that effector functions of antibodies are differentially induced and persist to different degrees across vaccine platforms during the primary vaccination series, and can be further tuned with boosting.

Previous work has shown that nearly all clinically approved COVID-19 vaccines elicit Spike-specific antibodies and can confer protection against wildtype virus induced disease (30, 39–42). The concentrations and functionalities of these antibodies vary across vaccine platforms (24, 28) and wane with time (20, 31, 43, 44). Here we observed that following the primary series of CoronaVac vaccination, complement depositing, opsinophagocytic and NK cell activating antibodies were observed. However, many of these functions were induced to a slightly lower level than mRNA vaccination, with limited breadth to Omicron. mRNA vaccination, conversely, drove robust antibody effector functions. Nevertheless, many of these functions waned over time, including a loss of complement deposition, neutrophil phagocytosis, and NK cell degranulation. In contrast, monocyte phagocytosis remained stable over 4-5 months following mRNA vaccination, pointing to limited-to-no waning of this critical antibody effector function. Therefore, despite the general loss of neutralization and antibody effector functions that may result in a renewed susceptibility to infection, the persistence of opsinophagocytic antibodies may continue to provide a first line of defense against infection, potentially providing some level of persistent protection against the virus months after mRNA vaccination. Thus, similar to opsinophagocytic mechanisms of protection against other mucosal pathogens (45), it is plausible that persistent protection against severe disease and death afforded by mRNA vaccination may be linked to this unique opsinophagocytic activity.

Despite the nearly complete loss of CoronaVac functional humoral immunity, months after the primary series, we still observed a robust rise in CoronaVac primed immunity following an mRNA boost. Strikingly, this boosting was observed even for antibody functions that were not induced robustly following the primary series. Moreover, mRNA boosting also expanded the breadth of the response, arguing that the heterologous boosting strategy resulted in functional maturation for both the antigen-binding (Fab)- and constant (Fc)-domain-associated antibody responses. Whether similar patterns of functional maturation of the humoral immune responses would be observed in the presence of alternate mix-and-match strategies remains unclear, but the data presented here strongly argue that combinations of these highly distributed vaccines could potentially be used in optimal combinations to drive broad pan-VOC functional immunity.

Analysis across VOCs highlighted significant differences in variant recognition by the FcR binding antibodies. Specifically, following the primary CoronaVac series, IgG1 antibodies exhibited some cross-VOC recognition; however, Coronavac-induced subclasses/isotypes and FcR binding antibodies exhibited little-to-no RBD and Spike recognition across Beta, Gamma, and Omicron. Only the CoronaVac-induced IgM responses exhibited cross VOC immunity. However, upon mRNA boosting, all antibody sucblass/isotypes/FcR binding increased to the WT spike, but subclass and FcR binding elevated to a lesser extent against the Gamma and Omicron Spike variants. Similarly, boosting drove robust FcR binding to the WT RBD, but more limited FcR binding to the Omicron RBD, despite strong binding antibodies to this variant antigen. Thus, these data suggest that binding to the RBD may not always be proportional to FcR binding and that other characteristics, including geometry and stoichemiotry, may also play an essential role in dictating antibody effector functions. Hence, further improvements in the quality of immunity may be achievable with future mix-and-match strategies, aimed at eliciting the strongest level of protective immunity against future VOCs.

Whether the heterologous nature of the mRNA boost or the timing of the delayed boosting drove improved CoronaVac Fab and Fc activity remains unclear. Yet, emerging data suggest that both the nature of the platform and the timing of immunization play a key role in the quality of the germinal center response, and thus humoral immune induction (46–49). Consequently, future studies are warranted to begin to probe all the mix-and-match and interval based effects of vaccine induced immune programming to help guide future rational vaccine campaigns aimed at driving the most protective vaccine induced immunity against SARS-CoV-2 and future variants.

The CoronaVac vaccine was the most widely distributed vaccine globally, with billions of doses administered worldwide (32, 47, 48). While the vaccine may not induce the most robust neutralizing antibody titers, protection against severe disease and death persisted across several VOCs (32). Moreover, this vaccine represents a unique priming strategy, given the exposure to all viral components, many of which may provide additional protection in the setting of future viral variation.

## Acknowledgments

We would like to thank the laboratory of Prof. Douglas Lauffenburger (Massachusetts Institute of Technology) for the critical evaluation of statistics in this manuscript. We also thank Mark and Lisa Schwartz, Terry and Susan Ragon, and the SAMANA Kay MGH Research Scholars award for their support. G.A. receives funding from the Massachusetts Consortium on Pathogen Readiness (MassCPR), the Gates Global Health Vaccine Accelerator Platform, and the NIH (3R37AI080289-11S1, R01AI146785, U19AI42790-01, U19AI135995-02, U19AI42790-01, P01AI1650721, U01CA260476 – 01, CIVIC75N93019C00052). We also would like to thank Estefany Poblete, Erick Salinas and Andres Muñoz for their excellent technical and professional expertise during clinical recruitment and sample processing. Work in the Medina laboratory was partially funded by the FONDECYT 1212023 grant from ANID of Chile, the FLUOMICS Consortium (NIH-NIAID grant U19AI135972) and the Center for Research on Influenza Pathogenesis (CRIP), an NIAID Center of Excellence for Influenza Research and Surveillance (CEIRS, contract # HHSN272201400008C). CPR and JL conducted this work as part of their Postdoctoral grant FONDECYT 3190706 and 3190648, respectively. MJA and ES conducted this work as part of their Ph.D. Thesis, under Programa de Doctorado en Ciencias Biológicas mención Genética Molecular y Microbiología, Facultad de Ciencias Biológicas, Pontificia Universidad Cátolica de Chile. MJA was funded by the ANID Becas/Doctorado Nacional 21212258 scholarship and ES was funded by a scholarship from Vicerrectoría de Investigación de la Escuela de Graduados, Pontificia Universidad Católica de Chile.

## Author Contributions

Conceptualization, X.T., R.P.M., M.J.A., R.A.M., and G.A.; Methodology, X.T., R.P.M., and G.A; Validation, X.T.; Formal Analysis, X.T., R.P.M., and G.A.; Investigation, X.T., R.P.M., G.A., M.J.A. and R.A.M.; Cohort study design, supervised and managed the sample collection, T.G.S., C.P.R., A.R. and R.A.M.; Processed samples, revised the paper, T.G.S., C.P.R., J.L., E.P., E.S., A.M., A.R. and R.A.M.; Resources, G.A. and R.A.M.; Writing – Original Draft, X.T. and R.P.M.; Writing – Review & Editing, R.P.M., G.A., M.J.A. and R.A.M.; Visualization, X.T., R.P.M., and G.A.; Project Administration, R.P.M.; Funding Acquisition, G.A. and R.A.M.; Supervision, R.P.M., R.A.M. and G.A.

## Methods

### Resource Availability

#### Lead contact

Further information and requests for resources and reagents should be directed to and will be fulfilled by the lead data point of contact, Ryan P. McNamara, or the corresponding authors, Galit Alter, or Rafael A. Medina.

#### Materials availability

This study did not generate new unique reagents.

#### Data and code availability

All coding was done using R Studio V 1.4.1103 using ggplot. Individual groups were analyzed as factors. Any additional information required to reanalyze the data reported in this paper is available from the lead contact upon request. All codes and scripts are available upon request to the lead data point of contact. The paper does not report original code.

### Experimental Model and Subject Details

Serum samples were obtained from subjects who received the complete-dosage regimen of the perspective vaccines recommended by the manufacturers. The cohort contains samples from individuals who received either BNT162b2 (n = 15) or CoronaVac vaccines (n = 34). The BNT162b2 mRNA vaccine group was given 30 μg BNT162b2 (23 – 53 years old, median: 36 years, 81% female) on days 0 and 21, and serum samples were taken up to 209 days after the second dose. The CoronaVac group (23 – 46 years old, median: 31 years old, 61% female) received two doses of 600 U CoronaVac four weeks apart and individuals were sampled 1-209 days after the second dose. A subgroup of CoronaVac subjects (n = 20) received a booster dose of the BNT162b2 mRNA vaccine and was sampled 14-19 days after the mRNA booster. We did not observe any immunocompromising comorbidities associated with the cohort.

### Method Details

#### Antigens

All antigens used in this study are listed in **Supplementary Table 1**. Most antigens were lyophilized powder and were resuspended in water to afford a final concentration of 0.5 mg/mL. Antigens that required biotinylation were treated with the NHS-Sulfo-LC-LC kit per the manufacturer’s instruction. Removal of excess biotin and buffer exchange from Tris-containing antigens was done using the Zebra-Spin desalting and size exclusion chromatography columns.

#### Immunoglobulin isotype and Fc receptor binding

Antigen-specific antibody levels of isotypes and subclasses and levels of Fcγ-receptor binding were evaluated using a custom multiplexing Luminex-based assay platform in technical replicates, as previously described (34). The antigens were directly coupled to magnetic Luminex beads (Luminex Corp, TX, USA) by carbodiimide-NHS ester-coupling chemistry, which designates each region to each antigen. Individual dilution curves for each antigen were performed to identify an appropriate dilution factor for each secondary feature that was within the linear range of detection. The antigen-coupled beads were incubated with different serum dilutions (1:100 for IgG2, IgG3, IgG4, IgM, and IgA1, 1:500 for IgG1, and 1:750 for Fcγ-receptor binding) overnight at 4°C in 384 well plates (Greiner Bio-One, Germany). Unbound antibodies were removed by washing and subclasses, isotypes were detected using the respective PE-conjugated antibody listed in **Supplementary Table 1**. All detection antibodies were used at a 1:100 dilution.

For the analysis of Fcγ-receptor binding PE-Streptavidin (Agilent Technologies, CA, USA) was coupled to recombinant and biotinylated human FcγR2A, FcγR2B, FcγR3AV, or FcγR3B protein. Coupled Fcγ-receptors were used as a secondary probe at a 1:1000 dilution. After 1 hour of incubation, the excessive secondary reagent was removed by washing, and the relative antibody concentration per antigen was determined on an IQue Screener PLUS cytometer (IntelliCyt).

#### Evaluation of antibody-mediated functions

Antibody-dependent cellular phagocytosis (ADCP) and neutrophil phagocytosis (ADNP) were evaluated using a flow cytometry-based phagocytic assay that requires the usage of fluorescently labeled microspheres, as described previously (34). The WT and Omicron Spike antigens were biotinylated and conjugated to yellow-green fluorescent neutravidin microspheres. Then diluted serum samples with the pre-determined concentrations (1:100) were incubated with the coupled antigens. The pre-formed immune complexes bound with microspheres were washed and incubated with a human monocyte cell line (THP-1) for ADCP function or with neutrophils collected from healthy donors’ blood samples to assess ADNP activity. Cells studied under ADNP assays were then stained with anti-CD66b Pac blue antibody to calculate the percentage of CD66b+ neutrophils. In both ADCP and ADNP assays cells were fixed with 4% paraformaldehyde (PFA) and identified by gating on single cells and microsphere-positive cells. Microsphere uptake was quantified as a phagocytosis score, calculated as the (percentage of microsphere-positive cells) x (MFI of microsphere-positive cells) divided by 100000.

Antibody-dependent complement deposition (ADCD) assessed the ability of antigen-specific 20 antibodies to bind complement component C3b, as described. Briefly, SARS-CoV-2 WT and Omicron Spike proteins were biotinylated, coupled to red fluorescent neutravidin microspheres, and then incubated with serum samples (dilution 1:40). Immune complexes were then washed, incubated with guinea pig complement, then washed with 15 mM EDTA. The level of complement deposition was measured by fluorescein-conjugated goat IgG that targets the guinea pig complement C3b and further analyzed by gating on the single microspheres and C3b+ events.

Antibody-dependent NK Cell activation (ADNKA) assessed the ability of antigen-specific antibodies to activate human NK cells to up-regulate the production of CD107a, IFN-γ, and CCL4 (MIP-1β), as described. Briefly, ELISA plates were coated with SARS-CoV-2 antigens. NK cells were isolated from buffy coats, obtained from healthy blood donors, using RosetteSep NK enrichment kit, and rested overnight with IL-15. Antigen-coated ELISA plates were then incubated with serum samples (dilution 1:10). NK cells were stained with anti-CD107a and treated with a protein-transport inhibitor and with brefeldin A to block degranulation. NK cells were then added to the immune complexes, labeled with surface staining antibodies including anti-CD3, anti-CD16, and anti-CD56, washed, fixed with 4% PFA, and permeabilized with PERM A/B to allow intracellular staining with anti-IFN-γ and anti-CCL4. NK cells were identified as CD3-CD56+ cells, and the level of NK cell activation was evaluated as CD107a+ IIFN-γ+ CCL4+.

### Quantification and Statistical Analysis

All data analysis was done using R Studio V 1.4.1103 or FlowJo. Statistical analysis was done using R studio or GraphPad Prism. No data point was omitted from the analysis. Box and whisker plots were generated using ggplot, calculating the mean and standard deviation for each factor. Technical replicates for each sample were performed and the mean between replicates was plotted.

### Additional Resources

An informed written consent form was obtained under protocol 200829003, which was reviewed and approved by the Health Sciences Scientific Ethics Committee at Pontificia Universidad Católica de Chile (PUC). This work was supervised and approved by the Mass General Institutional Review Board (IRB #2020P00955 and #2021P002628).

