## Supplementary Material for "Waning and boosting of functional humoral immunity to SARS-CoV-2"

### Supplementary Information

Includes Supplementary Titles and Legends for Supplementary Figures 1 – 3, Supplementary Figures 1 – 3, and Supplementary Table 1.

Supplementary Information Titles and Legends
Legends are also shown underneath figures

**Supplementary Figure 1. COVID-19 vaccines do not have off-target humoral activation.** (A) IgG1 specificity controls were quantified in the baseline (prior to immunization) (white, column 1), 1- and 2-dose BNT162b2 mRNA (blue, columns 2 and 3, waning periods in lanes 4 and 5), 1- and 2-dose CoronaVac (gray, columns 6 and 7, waning periods in lanes 8 and 9) via Luminex systems serology. Y-axis represents the mean fluorescence intensity (MFI) of a specific antigen. Shown are box and whiskers, along with individual data points, which represent the mean of technical replicates on individual patients. Antigens used were against human coronavirus OC-43 (HCoV-OC43) Spike, HCoV-HKU1 Spike, Influenza HA, and Ebola virus glycoprotein (EBOV GP). (B) Same as for A, but for FcγR2A, FcγR2B, FcγR3A, and FcγR3B binding for each antigen.

**Supplementary Figure 2. mRNA-vaccine boosting significantly enhances biophysical humoral recognition against VOC Spikes.** (A) Fold increases from 5-month wane windows (column 1) to post-mRNA-vaccine boost (column 2) for all VOC for FcRs are shown At the bottom right is the color legend for the VOC. Means were calculated using Luminex serology for each FcR for each VOC by timeframe. (B) Correlation heatmaps for WT Spike before and after mRNA-vaccine boost are shown; on the upper-right is the heatmap legend. On the right is a volcano plot of Pearson's Coefficients (x-axis) and p-values (y-axis) of pairwise antibody and Fcγ-receptor correlations of WT Spike in two-dose CoronaVac recipients (gray) and two-dose CoronaVac recipients boosted with an mRNA vaccine BNT162b2 (purple) showing a general trend towards a more tightly, and statistically significant, overall humoral response. (C – G) Same as (B), but for (C) Alpha VOC, (D) Beta VOC, (E) Gamma VOC, (F) Delta VOC, and (G) Omicron VOC.

**Supplementary Figure 3. mRNA-vaccine boosting restores waned macrophage inflammatory protein 1 beta (MIP-1b) expression in NK cells after incubation with WT Spike, and expands function to Omicron**. (Left) Antibody-dependent MIP-1b percent expression in NK cells was quantified in individuals at baseline (column 1), who received two doses of CoronaVac (columns 2 and 3), during the waning period (columns 4 and 5), and after mRNA-vaccine booster (column 6); WT Spike is shown in gray and Omicron is shown in purple. Peak functionality was set to responses after dose 2, and percent changes during the waning and boosting are shown above the timepoints. (Right) MIP-1b expression against WT Spike (gray) and Omicron Spike (blue) in individuals at baseline (column 1), after two doses of an mRNA vaccine (columns 2 and 3), and during the waning period (columns 4 and 5). Peak functionality was set to responses after dose 2, and percent changes during the waning and boosting are shown above the timepoints. Shown are the mean (solid line) and standard errors of the mean (SEM, shaded areas). * = $p < 0.05$, ** = $p < 0.01$ for all WT and Omicron comparisons (paired T-tests).

**Supplementary Figure 1**

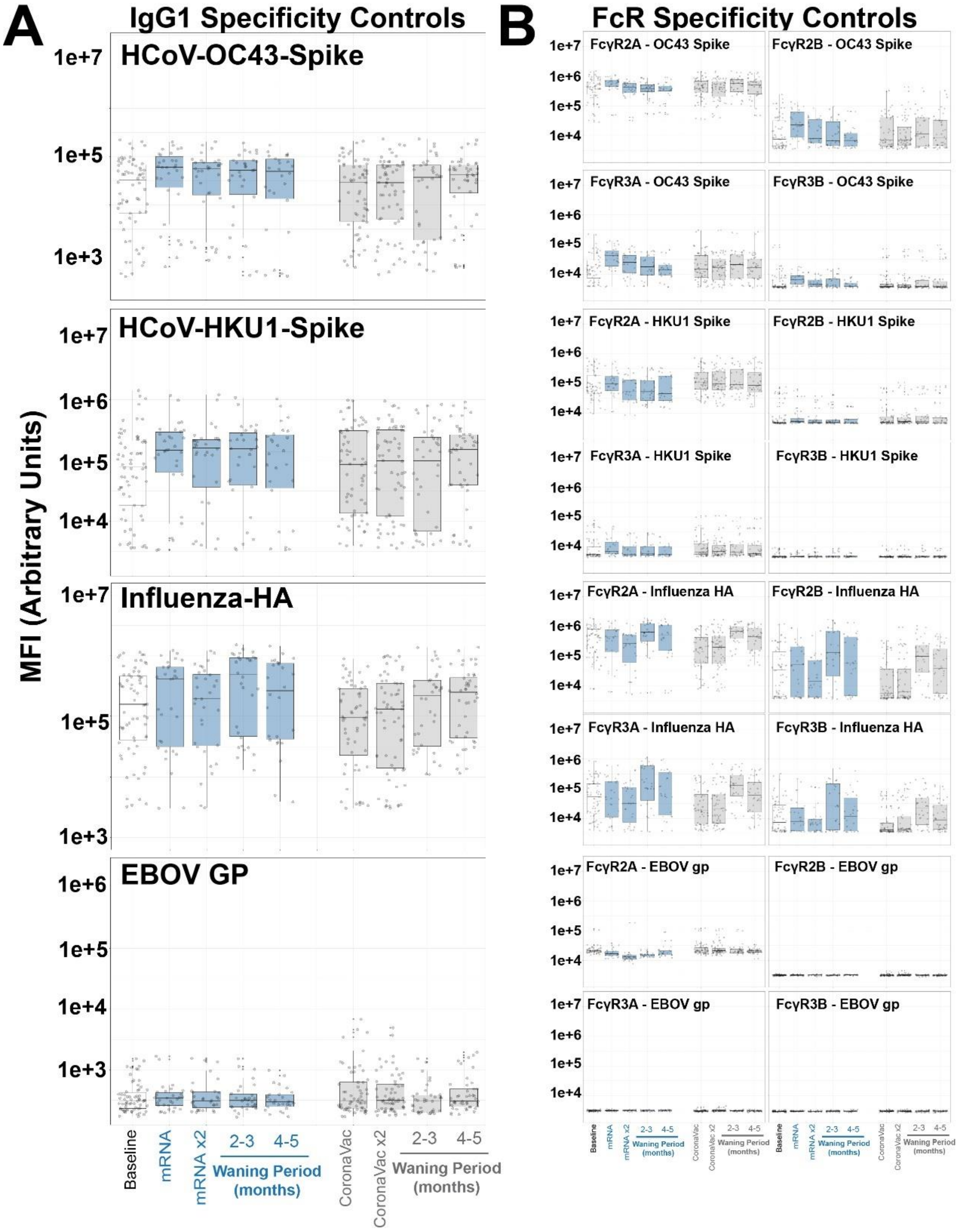


**Supplementary Figure 1. COVID-19 vaccines do not have off-target humoral activation.** (A) IgG1 specificity controls were quantified in the baseline (prior to immunization) (white, column 1), 1- and 2-dose BNT162b2 mRNA (blue, columns 2 and 3, waning periods in lanes 4 and 5), 1- and 2-dose CoronaVac (gray, columns 6 and 7, waning periods in lanes 8 and 9) via Luminex systems serology. Y-axis represents the mean fluorescence intensity (MFI) of a specific antigen. Shown are box and whiskers, along with individual data points, which represent the mean of technical replicates on individual patients. Antigens used were against human coronavirus OC-43 (HCoV-OC43) Spike, HCoV-HKU1 Spike, Influenza HA, and Ebola virus glycoprotein (EBOV GP). (B) Same as for A, but for FcγR2A, FcγR2B, FcγR3A, and FcγR3B binding for each antigen.

#### Supplementary Figure 2

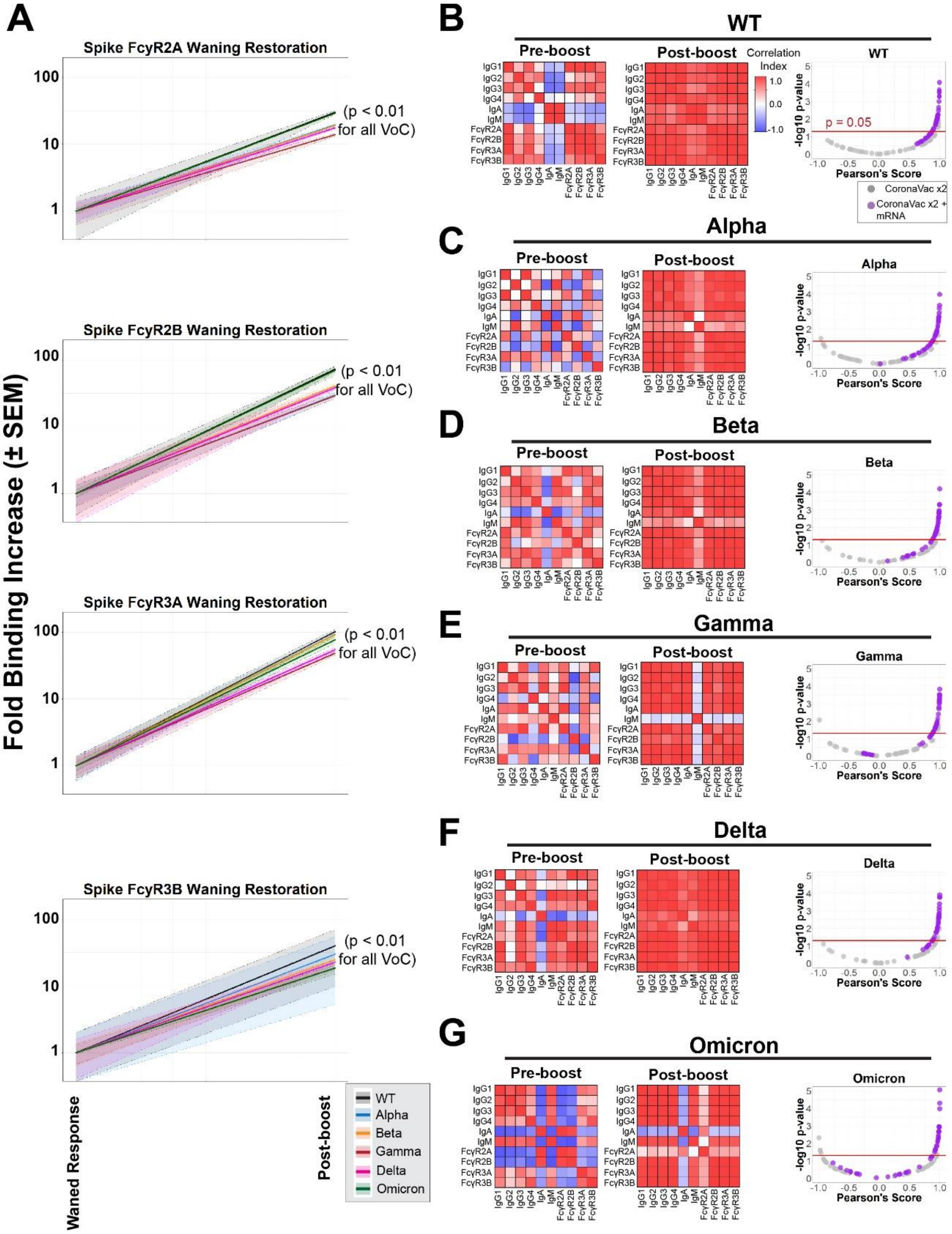


**Supplementary Figure 2. mRNA-vaccine boosting significantly enhances biophysical humoral recognition against VOC Spikes.** (A) Fold increases from 5-month wane windows (column 1) to post-mRNA-vaccine boost (column 2) for all VOC for FcRs are shown. At the bottom right is the color legend for the VOC. Means were calculated using Luminex serology for each FcR for each VOC by timeframe, and p-values were calculated from waned response to post-boost response. (B) Correlation heatmaps for WT Spike before and after mRNA-vaccine boost are shown; on the upper-right is the heatmap legend. On the right is a volcano plot of Pearson's Coefficients (x-axis) and p-values (y-axis) of pairwise antibody and Fcγ-receptor correlations of WT Spike in two-dose CoronaVac recipients (gray) and two-dose CoronaVac recipients boosted with an mRNA vaccine BNT162b2 (purple) showing a general trend towards a more tightly, and statistically significant, overall humoral response. (C – G) Same as (B), but for (C) Alpha VOC, (D) Beta VOC, (E) Gamma VOC, (F) Delta VOC, and (G) Omicron VOC.

**Supplementary Figure 3**

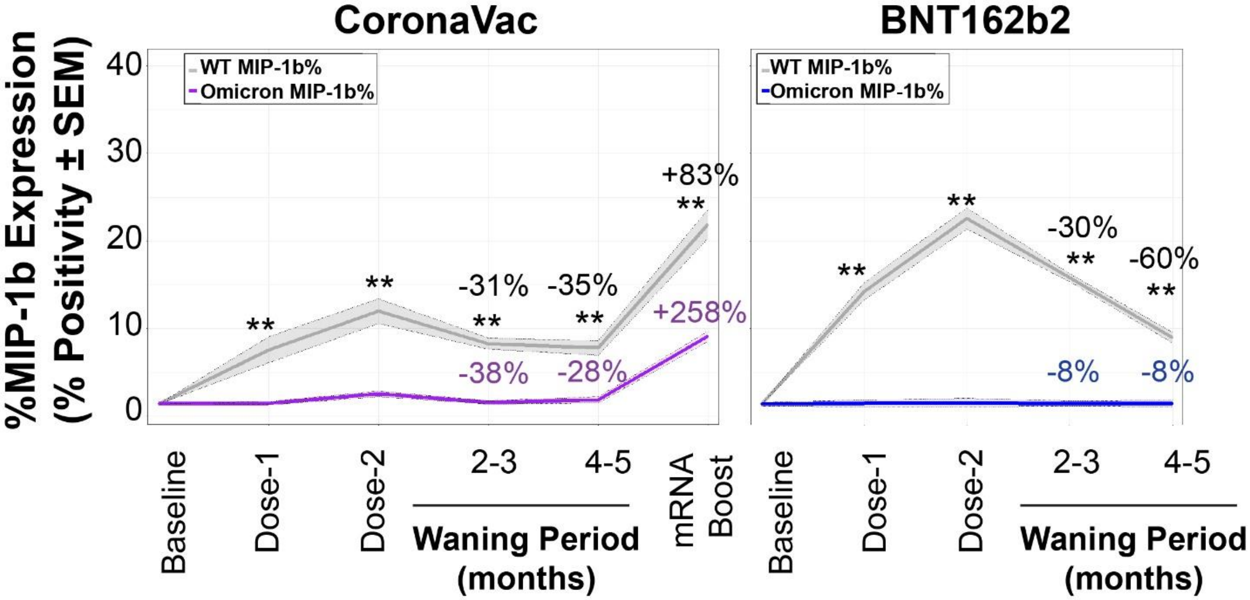


**Supplementary Figure 3. mRNA-vaccine boosting restores waned macrophage inflammatory protein 1 beta (MIP-1b) expression in NK cells after incubation with WT Spike, and expands function to Omicron**. (Left) Antibody-dependent MIP-1b percent expression in NK cells was quantified in individuals at baseline (column 1), who received two doses of CoronaVac (columns 2 and 3), during the waning period (columns 4 and 5), and after mRNA-vaccine booster (column 6); WT Spike is shown in gray and Omicron is shown in purple. Peak functionality was set to responses after dose 2, and percent changes during the waning and boosting are shown above the timepoints. (Right) MIP-1b expression against WT Spike (gray) and Omicron Spike (blue) in individuals at baseline (column 1), after two doses of an mRNA vaccine (columns 2 and 3), and during the waning period (columns 4 and 5). Peak functionality was set to responses after dose 2, and percent changes during the waning and boosting are shown above the timepoints. Shown are the mean (solid line) and standard errors of the mean (SEM, shaded areas). * = $p < 0.05$, ** = $p < 0.01$ for all WT and Omicron comparisons (paired T-tests).

#### Supplementary Table 1

| REAGENT or RESOURCE | SOURCE | IDENTIFIER |
|---|---|---|
| **Anti-Human IgG1-PE** | Southern Biotech | HP6001 |
| **Anti-human IgG2-PE** | Southern Biotech | 31-7-4 |
| **Anti-human IgG3-PE** | Southern Biotech | HP6050 |
| **Anti-human IgG4-PE** | Southern Biotech | HP6025 |
| **Anti-human IgM-PE** | Southern Biotech | SA-DA4 |
| **Anti-human IgA1-PE** | Southern Biotech | HP6025 |
| **Anti-CD66b Pac Blue** | BioLegend | 305112 |
| **Anti-CD107a** | BD Biosciences | 555802 |
| **Anti-CD3** | BD Biosciences | 558117 |
| **Anti-CD16** | BD Biosciences | 557758 |
| **Anti-CD56** | BD Biosciences | 557747 |
| **Anti-IFNγ** | BD Biosciences | 340449 |
| **Anti-CCL4** | BD Biosciences | 550078 |
| **Anti-C3b** | MP Biomed | 855385 |
| **SARS-CoV-2 WT Spike** | Sino Biological | 40589-V08H4 |
| **SARS-CoV-2 WT S1 Domain** | Sino Biological | 40591-V08H |
| **SARS-CoV-2 WT Receptor Binding Domain (RBD)** | Sino Biological | 40592-V08H |
| **SARS-CoV-2 WT S2 Domain** | Sino Biological | 40590-V08B |
| **SARS-CoV-2 WT N-terminal Domain** | Sino Biological | 40591-V49H |
| **SARS-CoV-2 Alpha Variant S** | Sino Biological | 40589-V08B6 |
| **SARS-CoV-2 Alpha Variant RBD** | Sino Biological | 40592-V08H82 |
| **SARS-CoV-2 Beta Variant S** | Sino Biological | 40589-V08B7 |
| **SARS-CoV-2 Beta Variant RBD** | Sino Biological | 40592-V08H59 |
| **SARS-CoV-2 Gamma Variant S** | Sino Biological | 40589-V08B10 |
| **SARS-CoV-2 Gamma Variant RBD** | Sino Biological | 40592-V08H86 |
| **SARS-CoV-2 Delta Variant S** | Sino Biological | 40589-V08B16 |
| **SARS-CoV-2 Delta Variant RBD** | Sino Biological | 40592-V08H115 |
| **SARS-CoV-2 Omicron Variant S** | Sino Biological | 40589-V08H26 |
| **SARS-CoV-2 Omicron Variant RBD** | Sino Biological | 40592-V08H121 |
| **Human Coronavirus OC43 S** | Sino Biological | 40607-V08B |
| **Human CoV HKU1 S (isolate N5)** | Sino Biological | 40606-V08B |
| **Human Cytomegalovirus (HCMV) Glycoprotein B (gB)** | Sino Biological | 10202-V08H1 |
| Ebola Virus Glyoprotein | IBT Bioservices | 0501-015 |
| **PE-Streptavidin** | Agilent Technologies | PB32-10 |
| **Guinea Pig Complement** | Cedarlane | CL4051 |
| **Protein Transport Inhibitor** | BD Biosciences | 554724 |
| **Brefeldin-A** | Sigma | B7651 |
| NHS-Sulfo-LC-LC Kit | ThermoFisher | 21435 |
| Zebra-Spin Desalting and Chromatography Columns | ThermoFisher | 89882 |
| RosetteSep NK Enrichment Kit | Stem Cell Technologies | 15065 |
| Fix & Perm Cell Permeabilization Kit | ThermoFisher | GAS002S-100 |
| R Studio V 1.4.1103 | RStudio, PBC | Open Source |
| GraphPad Prism | GraphPad Software, LLC | Ragon Site License |
| FlowJo V. 10.8 | FlowJo, LLC | www.flowjo.com/solutions/flowjo/downloads |

| | | |
|---|---|---|
| iQue Forecyt | Sartorius | 60028 |
| iQue Screener Plus | Intellicyt/Sartorius | 11811 |
| 384-well HydroSpeed Plate Washer | Tecan | 30190112 |
| MagPlex Microspheres | Luminex MFG | MC12001-01 (Cataloged by region) |
| Green Fluorescent Neutravidin Microspheres | ThermoFisher | F8776 |
| Red Fluorescent Neutravidin Microspheres | ThermoFisher | F8775 |

**Supplementary Table 1. List of reagents and resources used in this study.**
